# Massive expansion of human gut bacteriophage diversity

**DOI:** 10.1101/2020.09.03.280214

**Authors:** Luis F. Camarillo-Guerrero, Alexandre Almeida, Guillermo Rangel-Pineros, Robert D. Finn, Trevor D. Lawley

**Affiliations:** Host-Microbiota Interactions Laboratory, Wellcome Sanger Institute, Wellcome Genome Campus, Hinxton, CB10 1SA, UK; European Bioinformatics Institute (EMBL-EBI), Wellcome Genome Campus, Hinxton, CB10 1SA, UK; Wellcome Sanger Institute, Wellcome Genome Campus, Hinxton, CB10 1SA, UK

**Author notes:** Correspondence (L.F.C.), (T.D.L.).

## Abstract

Bacteriophages drive evolutionary change in bacterial communities by creating gene flow networks that fuel ecological adaptions. However, the extent of viral diversity and prevalence in the human gut remains largely unknown. Here, we introduce the Gut Phage Database (GPD), a collection of ∼142,000 non-redundant viral genomes (>10 kb) obtained by mining a dataset of 28,060 globally distributed human gut metagenomes and 2,898 reference genomes of cultured gut bacteria. Host assignment revealed that viral diversity is highest in the Firmicutes phyla and that ∼36% of viral clusters (VCs) are not restricted to a single species, creating gene flow networks across phylogenetically distinct bacterial species. Epidemiological analysis uncovered 280 globally distributed VCs found in at least 5 continents and a highly prevalent novel phage clade with features reminiscent of p-crAssphage. This high-quality, large-scale catalogue of phage genomes will improve future virome studies and enable ecological and evolutionary analysis of human gut bacteriophages.

## INTRODUCTION

Viruses are the most numerous biological entities on Earth with an estimated population size of 10^31^ particles (Brüssow and Hendrix, 2002). Bacteriophages (or phages; viruses that infect bacteria and archaea) profoundly influence microbial communities by functioning as vectors of horizontal gene transfer (Jain et al., 1999), encoding accessory functions of benefit to host bacterial species (Harrison and Brockhurst, 2017), and promoting dynamic co-evolutionary interactions (Betts et al., 2014). For decades, the discovery of phages occurred at a slow pace. However, with the advent of high-throughput metagenomics, it became possible to uncover an unparalleled amount of novel phage diversity (Al-Shayeb et al., 2020; Paez-Espino et al., 2016). A surprising finding was that the majority of phage sequences uncovered by metagenomics could not be classified into any known viral taxonomy laid out by the International Committee on Taxonomy of Viruses (ICTV) (e.g. species, genus, family) (Simmonds et al., 2017) prompting many researchers to organize phage predictions from metagenomic datasets into grouping schemes based solely on genomic features (Bin Jang et al., 2019).

The impact of phages on different ecosystems is beginning to be uncovered, with phages found in the oceans already being referred to as ‘puppet masters’ due to their significant impact on oceanic biogeochemistry (Breitbart et al., 2018). Given the impact of the gut microbiome composition and function on human health, there is a growing focus on phages that inhabit the gut ecosystem (Clooney et al., 2019; Kho and Lal, 2018). The first metagenomic studies revealed that the majority (81%-93%) of the viral gut diversity is novel (Manrique et al., 2016; Reyes et al., 2010) but gut phage host assignment and host range remain largely uncharacterized. An exception has been p-crAssphage, a phage discovered in 2014 by computational analysis of metagenomic reads and found in >50% of Western human gut microbiomes (Dutilh et al., 2014). Analyses of predicted phage sequences from gut metagenomes have yielded fascinating insights into phage biology, such as the presence of sticky domains — which may facilitate adherence of phage to the intestinal mucus (Barr et al., 2013) — reverse transcriptases that promote gene hypervariation (Minot et al., 2012), and proteins with ankyrin domains that may aid bacterial hosts in immune evasion (Jahn et al., 2019)

Previous analyses have focused on bulk viral fragments with limited resolution to characterize individual phage genomes or link specific phage to a bacterial host species (Minot et al., 2012). More recently, human gut metagenomes have been mined to compile a more comprehensive list of gut phage genomes (Gregory et al., 2019; Paez-Espino et al., 2019), providing new fundamental insights into the viral diversity and functions present in the human gut microbiome. Nevertheless, the limited number (<700) of metagenomes used to construct these databases (GVD and gut phage fraction from IMG/VR), and the fragment size of their predictions (median size <15 kb as opposed to ∼50 kb for an average *Caudovirales* phage genome commonly found in the human gut), suggests that the majority of gut phage diversity remains uncharacterized and incomplete. Indeed, a recent report estimated that IMG/VR, which contains viral sequences from a wide range of environments, showed that only 1.9% of the predictions were complete, and 2.5% were classified as high-quality (>90% complete)(Nayfach et al., 2020). A comprehensive resource of longer and complete reference phage genomes is required to enable genome-resolved metagenomics for gut phage studies across human populations.

Here, we introduce the Gut Phage Database (GPD), a highly curated database containing 142,809 non-redundant phage genomes derived from the analysis of 28,060 globally distributed metagenomic samples. Importantly, the GPD includes over 40,000 high-quality genomes with a median size of 47.68 kb. We use GPD to gain insight into the biology, host range and global epidemiology of human gut phages. We uncover 280 globally distributed viral clusters, including one viral clade (Gubaphage) with reminiscent features to p-crAssphage. Given the high quality of the reference genomes, the database size, and the sequence diversity harboured by the GPD, this resource will greatly improve the characterization of individual human gut bacteriophages at a global or local scale.

## RESULTS

### Generation of the Gut Phage Database (GPD)

In order to obtain a comprehensive view of human gut phage diversity, we analysed 28,060 public human gut metagenomes and 2898 bacterial isolate genomes cultured from the human gut (Figure 1A). To identify viral sequences among human gut metagenomes, we screened over 45 million assembled contigs with VirFinder (Ren et al., 2017) which relies on *k*-mer signatures to discriminate viral from bacterial contigs, and VirSorter (Roux et al., 2015) which exploits sequence similarity to known phage and other viral-like features such as GC skew. Since obtaining high-quality genomes was essential for our downstream analyses, we used conservative settings (see “Methods” section for further details) for both tools and retained only predictions that were at least 10 kb long.

**Figure 1.**
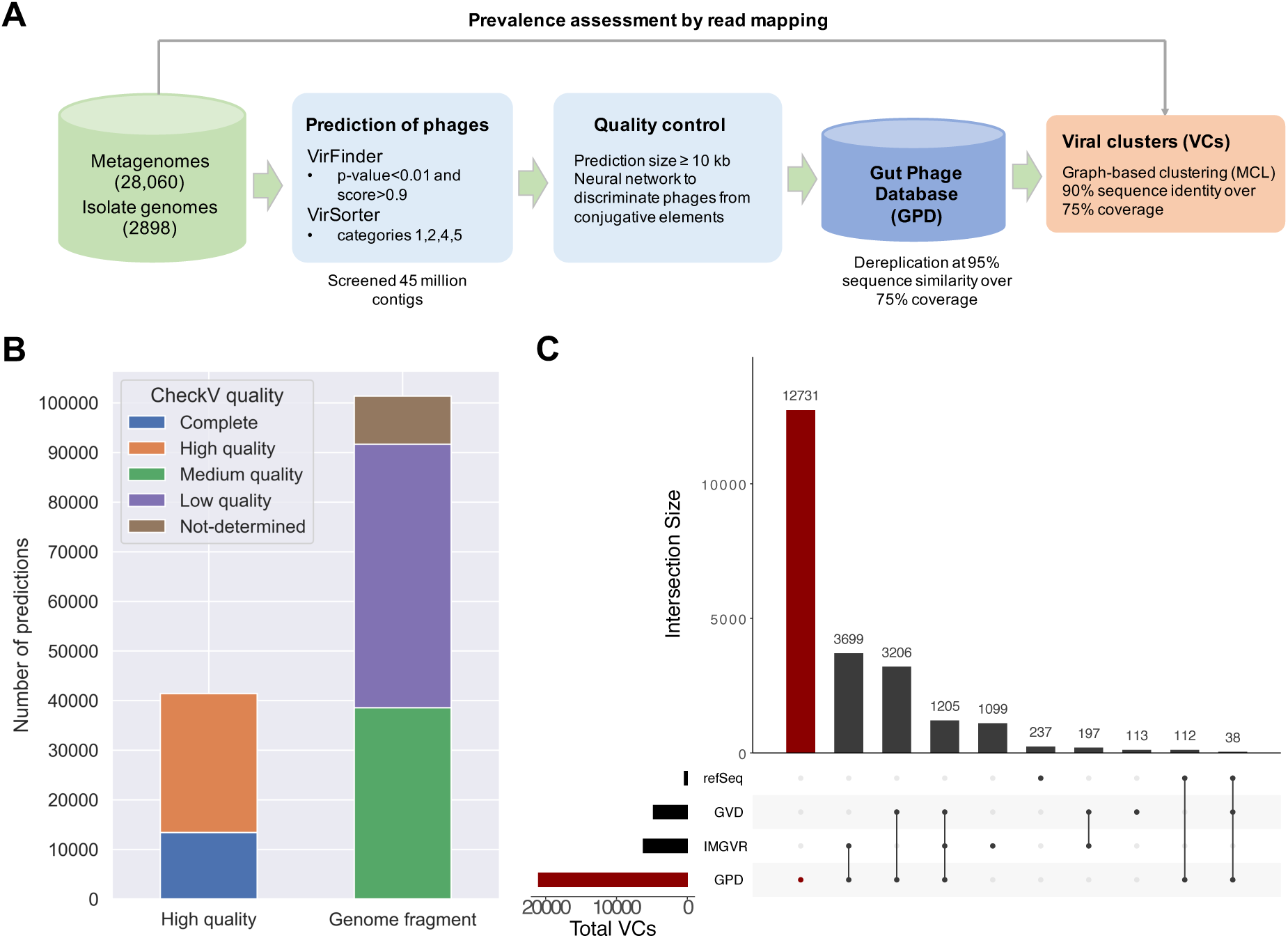
Generating the most complete sequence database of human gut bacteriophages. **A)** Massive prediction of phage genomes from 28,060 human gut metagenomes and 2898 isolate genomes was carried out using VirFinder and VirSorter with conservative settings. A machine learning approach (see “Methods”) was used to increase the quality of predictions and redundancy was removed by clustering the sequences at a 95% sequence identity. Diversity was further analysed by generating viral clusters (VCs) of predictions using a graph-based approach. **B)** Quality estimation of GPD genomes by CheckV. Over 40,000 predictions are categorized as high-quality. **C)** UpSet plot comparing GPD against other public gut phage databases. GPD captures the greatest unique diversity of phage genomes that inhabit the human gut.

To further improve the quality of the dataset, we devised a machine learning approach to filter out contaminant mobile genetic elements (Figure S1A). We identified predictions carrying machinery from type IV secretion systems, suggesting contamination by integrative and conjugative elements (ICEs). We used a feedforward neural network to discriminate phages from ICEs by exploiting differences in gene density, fraction of hypothetical proteins, and k-mer composition signatures (see “Methods” section). The classifier was trained with experimental sequences of phages and ICEs and showed an excellent performance in an independent test set (AUC>0.97) (Figure S1B) of human gut mobile genetic elements (MGEs). Next, we dereplicated the final set of filtered sequences at a 95% average nucleotide identity (ANI) threshold (over a 75% aligned fraction) obtaining a database of 142,809 gut phage sequences, henceforth referred to as the Gut Phage Database (GPD).

We estimated the level of completeness of each viral genome using CheckV (Nayfach et al., 2020) (Figure 2B). This tool infers the expected genome length of a viral prediction based on the average amino acid identity to a database of complete viral genomes from NCBI and environmental samples. In total, 13,429 (9.4%) of the viral genomes were classified as complete, 27,999 (19.6%) as high-quality, and 101,381 (70.99%) as genome fragments (<90% complete). This classification scheme is consistent with the MIUViG standards (Roux et al., 2019). The median genome completeness of all genomes stored in the GPD was estimated to be 63.5% (interquartile range, IQR= 34.68%–95.31%) (Figure S1C). Estimation of non-viral DNA by CheckV showed that 73.5% of GPD predictions had no contamination whereas 84.13% had a predicted contamination <10% (Figure S2D). In comparison to other human gut phage databases (Gregory et al., 2019; Paez-Espino et al., 2019), GPD had the largest median genome size with ∼31 kb, followed by IMG/VR and GVD with 15 kb and 11 kb, respectively (Figure S1E).

**Figure 2.**
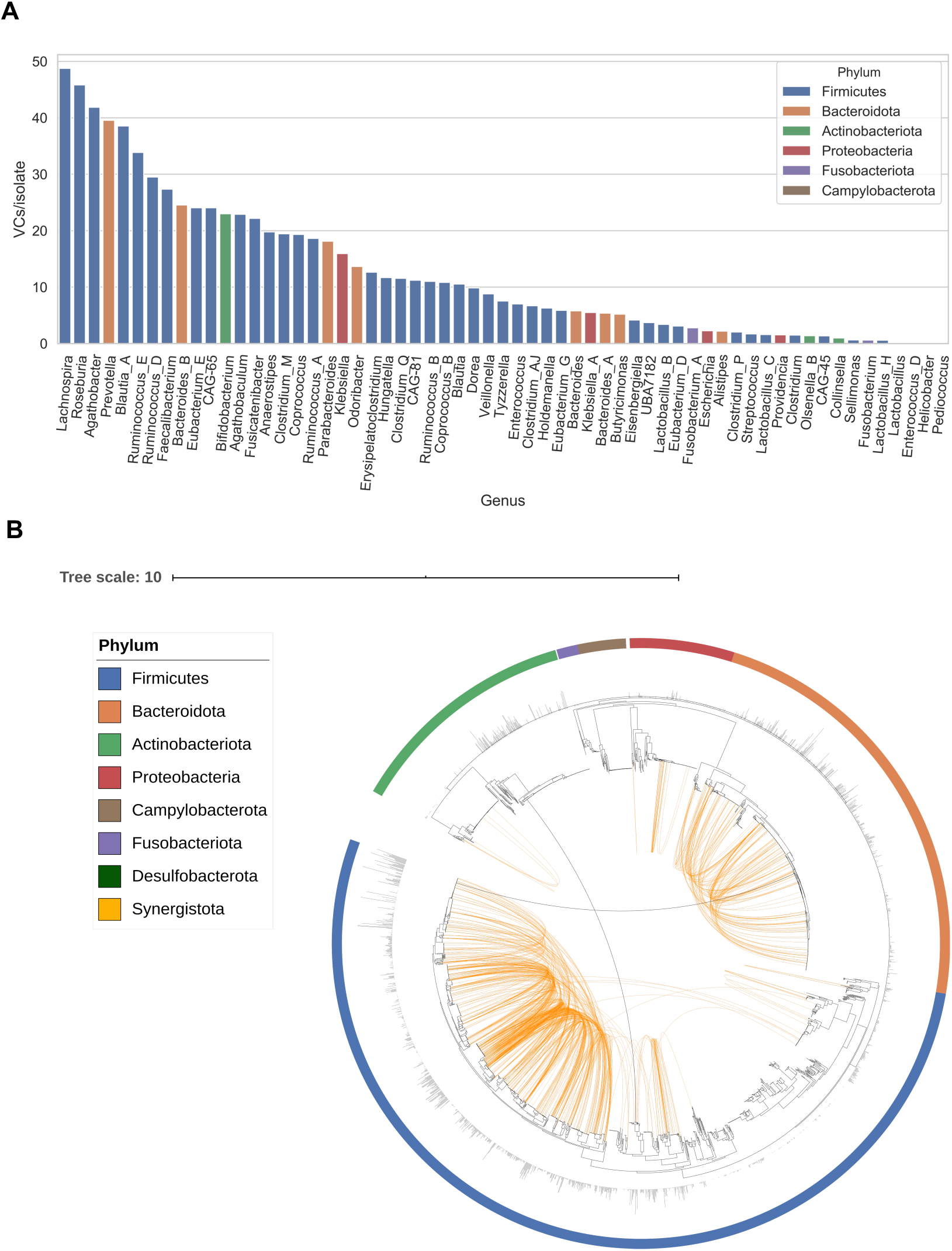
Bacterial host assignment and host range for gut phage. **A)** Bacterial genera with the highest viral diversity were *Lachnospira, Roseburia, Agathobacter, Prevotella*, and *Blautia* A. On the other hand, the lowest viral diversity was harboured by *Helicobacter* and the lactic acid bacteria *Lactobacillus*, Lactobacillus *H, Enterococcus* D and *Pediococcus*. **B)** Phylogenetic tree of 2898 gut bacteria isolates showing phage host range. Host assignment was carried out by linking prophages with their assemblies and CRISPR spacer matching. Orange connections represent VCs with a very broad host range (not restricted to a single genus). Black connections represent VCs able to infect two phyla. Outer bars show phage diversity (VCs/isolate).

### GPD significantly expands gut bacteriophage diversity

In order to assess the viral diversity of the GPD at high taxonomic levels, we used a graph-based clustering approach to group genetically related phages. Merging GPD with the RefSeq phages and two other human gut phage databases (GVD and gut phage fraction of IMG/VR) resulted in the generation of 21,012 non-singleton viral clusters (VCs) with at least 1 GPD prediction (GPD VCs). A VC corresponds to a viral population sharing approximately 90% sequence identity over ∼75% aligned fraction (see “Methods” section for further details). Benchmarking against the RefSeq phages (Brister et al., 2015) revealed that the boundaries of GPD VCs were equivalent to a subgenus level, as 99.73% of all VCs were contained within the genus level.

Strikingly, less than 1% (171 out of 21,012) of the GPD VCs overlap with the RefSeq phages. Phages from these 171 VCs mainly infect *Escherichia, Enterobacter, Staphylococcus*, and *Klebsiella* genera, reflecting the bias of the RefSeq database towards well-known clinically important and traditionally cultured bacteria. Consistent with previous reports of phage predictions from metagenomic datasets (Hoyles et al., 2014), we were not able to confidently assign a family to the majority (∼80%) of GPD VCs, while the rest corresponded mainly to the *Podoviridae, Siphoviridae* and *Myoviridae* families (Figure S1E). These 3 viral families belong to the *Caudovirales* order (phages characterized by having tails and icosahedral capsids) which were previously reported to be enriched in human faeces (Hoyles et al., 2014; Roux et al., 2012).

For comparison purposes, we also considered VCs without GPD predictions (Figure 1C). Analysis of VCs composed from only GPD and IMG/VR genomes showed 3,699 overlaps, while we found 3,206 VCs composed of only GPD and GVD genomes. Moreover, GPD harboured the highest number of unique VCs with 12,731 novel clusters. On the other hand, 1099 VCs and 113 VCs were unique to IMG/VR and GVD, respectively. In addition, 1205 VCs were shared by the three databases. Interestingly, the number of VCs with an assigned phage taxon was lower in the VCs that were unique to GPD as opposed to those shared with GVD and IMG/VR (18.74% vs 27.8%) (*P=*1.96⨯ 10^−9^, *χ*^2^ test). Thus, GPD considerably expanded the previously unknown gut phage diversity in the human gut. This phage diversity expansion is likely driven by the high number of gut metagenomes mined and their global distribution which allows the retrieval of rarer gut phage clades.

### Bacterial host assignment and host range for gut phage

The GPD creates a unique opportunity to assign specific phage to bacterial host species at an unprecedented scale providing a phylogenetic framework to study gut bacteria-phage biology. Accordingly, we inferred the most likely bacterial hosts for each phage prediction using a comprehensive collection of 2898 high-quality human gut bacterial isolate genomes (Forster et al., 2019; Zou et al., 2019). By screening for the presence of CRISPR spacers targeting phage and by linking the prophages to their assemblies of origin (Edwards et al., 2016), we assigned 40,932 GPD phage (28.66% of all predictions) to 2,157 host strains. This corresponded to at least one phage for 74.43% of all cultured human gut bacteria. We then analysed if there was any preference for phage infection across 4 common human gut bacterial phyla (Firmicutes, Bacteroidetes, Proteobacteria, and Actinobacteriota) (Figure S2A). At the phylum level, we detected significant lower phage prevalence in Actinobacteriota, with 58.79% infected isolates compared to at least 70% for the other phyla.

We then measured viral diversity (measured by the number of VCs per isolate) within each phylum. This analysis revealed that the Firmicutes harbour a significantly higher viral diversity (Figure S2B), with an average of 3.13 VCs/isolate while also harbouring 60% of the total VCs assigned across all phyla. Interestingly, the Firmicutes diversity was unevenly distributed as most of the viral diversity originated from the Negativicutes and Clostridia classes, with an average of 4.88 VCs and 3.9 VCs per isolate respectively in contrast with the Bacilli (0.99 VC/isolate), and none for Bacilli A and Desulfitobacteriia classes.

Analysis at the bacterial genus level across all phyla revealed that *Lachnospira, Roseburia, Agathobacte*r, *Prevotella*, and *Blautia* A contain the highest number of VCs/isolate (Figure 2A). With the exception of *Prevotella*, which belongs to the Gram-negative Prevotellaceae family, these genera are members of the Gram-positive Lachnospiraceae family of Firmicutes associated with butyrate-producing spore-formers. In contrast, the lowest viral diversity per isolate was detected among *Helicobacter*, and the lactic acid bacteria *Lactobacillus* H, *Lactobacillus, Enterococcus* D and *Pediococcus*. Thus, we observe a wide distribution of phage abundance and prevalence across human gut bacteria, even within the same phylum.

Horizontal transfer of genes between bacteria via transduction is a major driver of gene flow in bacterial communities (Chen et al., 2018). Host tropism of bacteriophage is believed to be limited by phylogenetic barriers, with most phages being usually restricted to a single host bacterial species (Ackermann, 1998). However, this has not been investigated at large scale across the human gut bacteria. Host assignment at different bacterial taxonomic ranks revealed that the majority of VCs were restricted to infect a single species (64.51%) (Figure S2C). We also found many VCs with broader host ranges such as those restricted to a single genus (22.39%), family (10.79%), order (1.86%), class (0.26%) and phylum (0.13%). Our findings are in line with a recent survey of the host range of gut phages by meta3C proximity ligation (6,651 unique host-phage pairs) which found that ∼69% of gut phages were restricted to a single species (Marbouty et al., 2020). Visualization of very broad range VCs (i.e. those not restricted to a single genus) reveals the large-scale connectivity between phylogenetically distinct bacterial species that fuels bacteria adaptation and evolution (Figure 2B). In general, the higher the viral diversity per bacterial genus, the higher the number of phages with broad host range (Figure S2D).

Surprisingly, two VCs (VC_269 and VC_644) had a host range that spanned two bacterial phyla. VC_269 was predicted to infect *Faecalibacterium prausnitzii* C (Firmicutes) and two *Bifidobacterium spp*. (Actinobacteriota), while VC_644 had a host range that included 5 *Bacteroides spp*. (Bacteroidota) and *Blautia* A *wexlerae* (Firmicutes). We predicted VC_269 to be a *Myoviridae* phage, on the other hand, we could not assign a taxonomy rank to VC_644. The presence of integrases in both VCs suggest that these are temperate phages. We hypothesize that additional phages infecting both Actinobacteriota and Firmicutes may be more common, as recent evidence supports a shared ancestry between phages that infect both Actinobacteria (*Streptomyces*) and Firmicutes (*Faecalibacterium*) (Koert et al., 2019).

Taken together, we reveal that approximately one third of gut phage have a broad host range not limited to a single host species. Our analysis provides a comprehensive blueprint of phage mediated gene flow networks in human gut microbiome.

### Human lifestyle associated with global gut distribution of phageome types

The gut phageome can be defined as the aggregate of phages that inhabit the gut (Manrique et al., 2016). We performed the most comprehensive phageome profiling of the human gut by read mapping 28,060 metagenomes against the GPD. These metagenomic datasets used to generate the GPD were sampled from 28 different countries across the six major continents (Africa, Asia, Europe, North America, South America and Oceania). Our initial analysis demonstrated a positive correlation between sample sequencing depth and the number of viral genomes detected for samples with <50 million reads. Therefore, we focused further analysis on a dataset of 3011 deeply sequenced (>50 million reads) metagenome samples spanning all continents and 23 countries (Figure S3A).

We observed clear separation of the North American, European, and Asian phageomes from African and South American samples when we computed the inter-sample Jaccard distance (Figure 3A) (*P =* 0.001, PERMANOVA test). Interestingly, these phageome patterns are associated with important differences in human lifestyles. Country-wise, samples derived from Africa and South America were mainly sampled from Peru, Tanzania, and Madagascar. Specifically, Peruvian and Tanzanian samples originate from hunter gatherer communities whereas Malagasy samples come from rural communities with non-Western lifestyles. Oceania was a special case because it had a similar fraction of samples belonging to both groups. However, when we stratified by country, we revealed that all Fijian samples clustered with the rural group, whereas Australian samples segregated with the urbanized cluster. Fiji samples were derived from rural agrarian communities. These observations support the hypothesis that lifestyle, particularly urbanization, may drive differences in the gut phageomes across different human populations.

**Figure 3.**
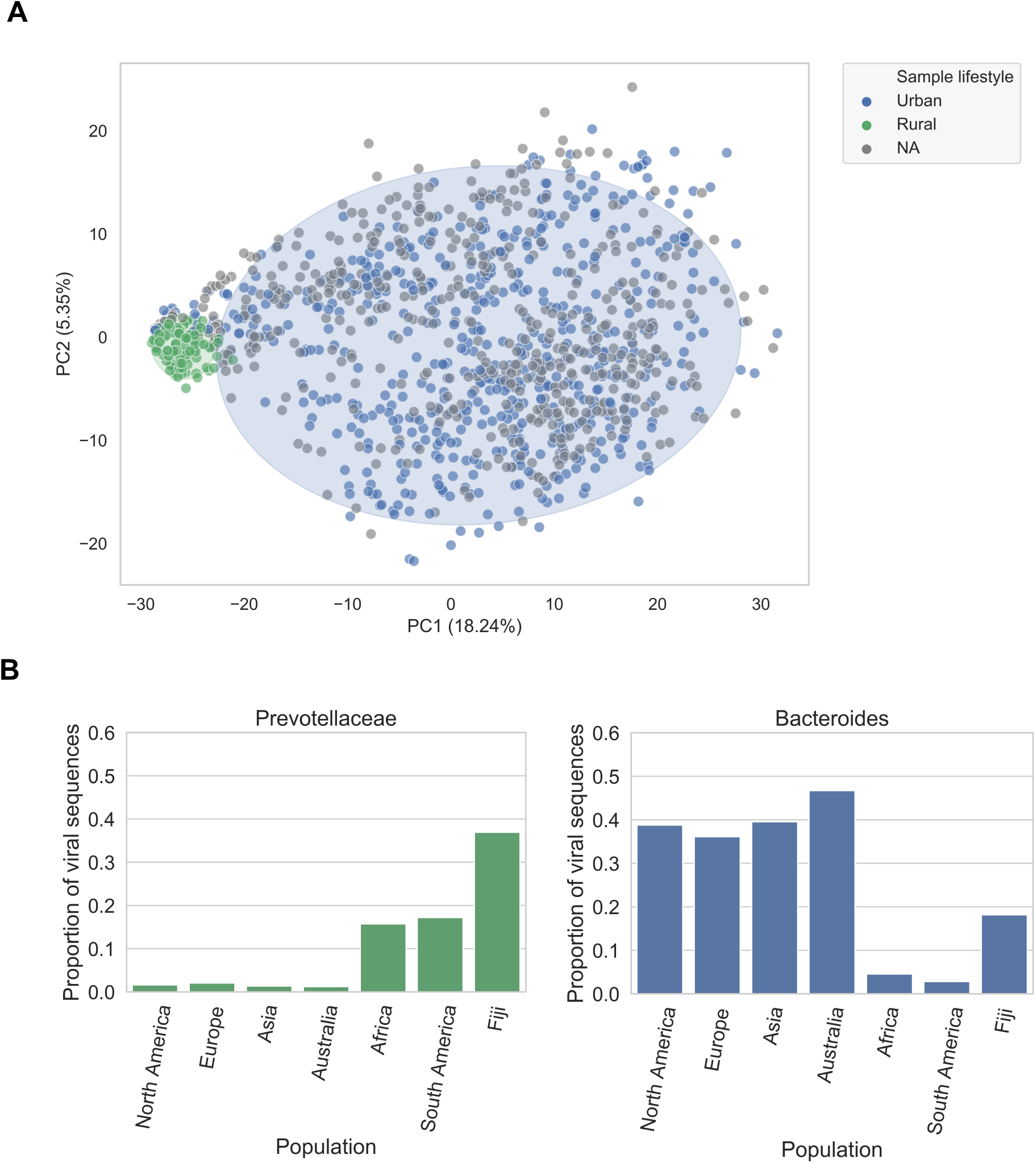
Global phylogeography of gut phages. **A)** PCA plot of inter-sample Jaccard distance. Lifestyle is associated with differences in the gut phageome across human populations. Samples from Peru, Madagascar, Tanzania and Fiji are found in the rural cluster whereas those samples with a more Westernized lifestyle (mainly from North America, Europe, and Asia) are found in the urban cluster (*P =* 0.001, R^2^ = 0.36, PERMANOVA test). Ellipses enclose samples within 2 standard deviations for each lifestyle. **B)** The proportion of viral sequences (at 95% sequence identity dereplicated) that target Prevotellaceae hosts in traditional societies is higher than that of industrialized populations. Conversely, Bacteroides hosts are more common in industrialized populations than in traditional societies. This result suggests that the composition of the gut phageome at a global scale is driven by the bacterial composition.

We reasoned that the bacterial composition of an individual’s microbiome would shape the gut phageome. Prevotellaceae bacteria are more abundant and prevalent in individuals living a rural/traditional lifestyle, whereas *Bacteroides* are more abundant and prevalent in individuals living a urban/Western lifestyle (Wu et al., 2011). By harnessing the host assignment data for each phage, we found a significantly higher proportion of VCs assigned to the Prevotellaceae family from African, South American and Fijian metagenome samples than that of North America, Europe, Asia, and Australia (*P* = 0.0, *χ*^2^ test) (Figure 3B). Conversely, the *Bacteroides* phages were significantly more prevalent in North America, Europe, Asia, and Australia gut microbiomes (*P* = 1.72⨯ 10^−208^, *χ*^2^ test). Given the correlation between microbiome enterotypes and phageome types, driven by the intimate connection between phages and their bacterial hosts, we provide evidence that human lifestyle is associated with global variation of gut phageomes, most likely mediated by differences in the host gut microbiome.

### Global distribution of 280 dominant human gut phages

If the gut phageome is predominantly shaped by the bacterial composition, we would expect to observe strong correlation between the prevalence of VCs with that of their bacterial hosts. A clear example is the crass-like family of gut phages which can be divided into 10 phage genera (Guerin et al., 2018). Genus I, which has been found in a large fraction of Western microbiome samples is able to infect species from the *Bacteroides* genus. In contrast, genera VI, VIII and IV were previously found to be the most prevalent crass-like phage among Malawian samples (Guerin et al., 2018). Here, we predict that the most probable host of these three phage genera is *Prevotella copri* (rest of crAss-like family predicted hosts in Table S1). In accordance with the results from the Malawian samples, we also found the prevalence of genera VI, VIII and IV to be higher than genus I in Africa, South America, and Fiji (Figure 4A). Thus, the crass-like family is globally distributed with distinct global distribution patterns at the genera level, which appears to be strongly influenced by human lifestyles and enterotypes.

**Figure 4.**
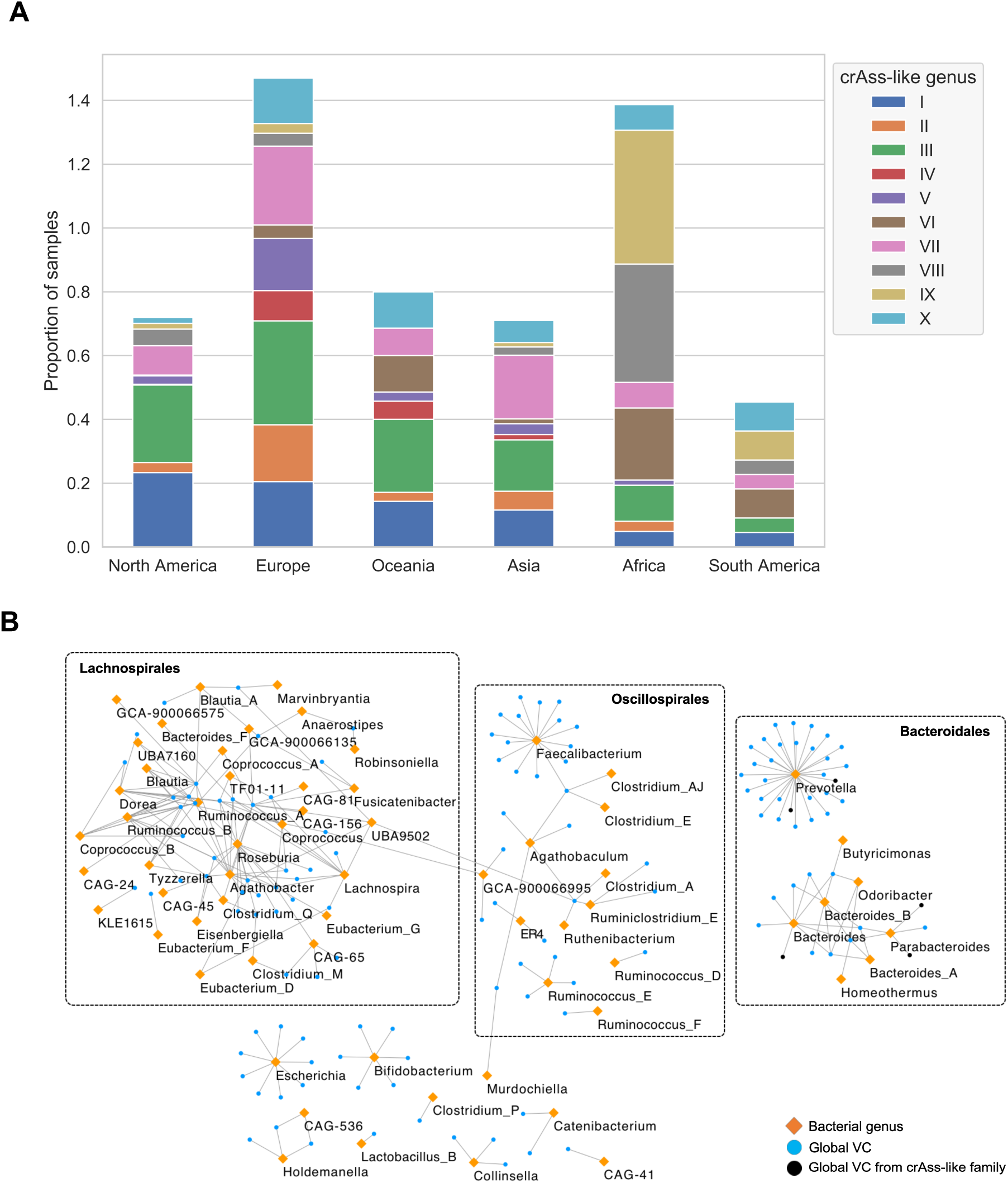
Global gut phage clades and their bacterial hosts. **A)** The crass-like family is a globally distributed phage. Genera VI, VIII and IX which are predicted to infect a *Prevotella* host are more common in Africa and South America in contrast to genus I which infects a *Bacteroides* host. **B)** Host-phage network of globally distributed VCs (orange) reveals that *Prevotella, Faecalibacterium*, and *Roseburia* are the most targeted bacterial genera. In contrast to the Bacteroidales and Oscillospirales, the VCs from the Lachnospirales are highly shared. VCs that belong to the crAss-like family are highlighted in black; These were predicted to infect *Prevotella, Bacteroides*, and *Parabacteroides*.

We further investigated if we could identify other gut phage VCs with global distributions. By extending the analysis to all the VCs we were able to detect a total of 280 VCs that were globally distributed (found in at least 5 continents). This represents ∼1.3% of all defined VCs (280/21,012). For 119 out of the 280 VCs (42.5%), we were able to classify them to the *Caudovirales* order, whereas the remaining 57.5% remained unclassified. Thus, the majority of globally distributed VCs are completely novel. When we looked at viral families detected within the *Caudovirales*, we detected *Podoviridae* (10 VCs), *Myoviridae* (28 VCs), *Siphoviridae* (43 VCs), and the newly formed family *Herelleviridae* (1 VC). In addition, when we examined at the phage subfamily level, the most common hits corresponded to the *Picovirinae* and *Peduovirinae* subfamilies with 4 VCs each. Importantly, the genomes of 131 members of 57 globally distributed VCs were mined directly from genomes of cultured isolates, providing unique opportunities for follow-up experiments to study bacteria-phage co-evolution (Table S2).

A bacteria-phage network of globally distributed VCs (Figure 4B) revealed that *Prevotella* was the most targeted genus (37 VCs), followed by *Faecalibacterium* and *Roseburia* with 15 VCs each. In addition, we observed that in contrast to the Bacteroidales and Oscillospirales, the global VCs associated to the Lachnospirales were highly shared between different genera (Figure S4A). Notably, whilst 12 globally distributed VCs were members of the crAss-like family (in black), we were only able to assign a host to 6 VCs which targeted Bacteroidales bacteria. We observed that globally distributed phages had a significant broader range (across different genera) than phages found in single continents (*P* = 1.62⨯10^−5^) (Figure S4B). This result suggests that broad host-range of certain VCs likely contribute to their expansion across human populations.

Thus, we show that along with 12 crass-like VCs, there exists a set of at least 280 VCs which are globally distributed. Functional characterization of members of this set will prove useful to shed light on what makes a gut phage to become widespread across human populations.

### The Gubaphage is a novel and highly prevalent clade in the human gut

When we calculated the number of genomes per VC, we discovered that VC_3 had the highest number of GPD predictions, only surpassed by VC_1 (which was composed of p-crAssphage genomes) (Figure 5A). Similarly to p-crAssphage, VC_3 was characterized by a long genome (∼80kb), a BACON domain-containing protein, and predicted *Bacteroides* host range. Searching for sequences in the GPD with significant similarity to VC_3 large terminase gene (E-value<1e-6), we identified other 205 related VCs. We refer to this clade of phages as the Gut Bacteroidales phage (Gubaphage). Given its reminiscent features to crAssphage, we decided to investigate if the Gubaphage belonged to the recently proposed crAss-like family which consists of 10 genera and 4 subfamilies. We examined this relationship by building a phylogenetic tree using the large terminase gene (Figure S5). The tree successfully clustered all the crAss-like genera as expected, however the Gubaphage significantly diverged from the other crAss-like phages forming a distinct clade.

**Figure 5.**
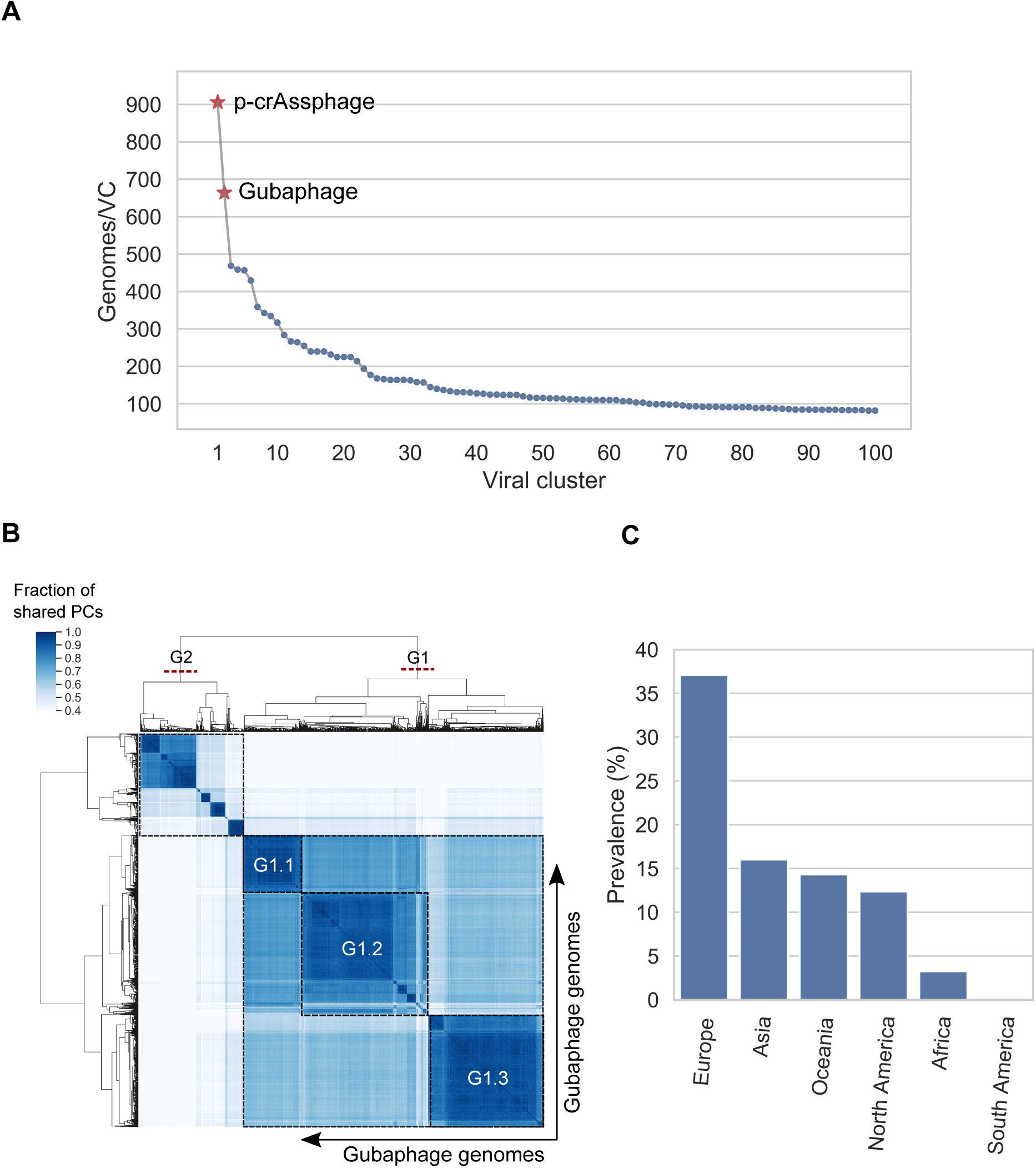
The Gubaphage is a novel and highly prevalent clade in the gut. **A)** VCs composed of only GPD predictions ranked by number of genomes. VC_3 which belongs to the Gubaphage clade was the second biggest cluster after VC_1 (composed of p-crAssphage genomes). **B)** Analysis of Gubaphage phylogenetic structure revealed two genera infecting members of the *Bacteroides* (G1) and *Parabacteroides* (G2). **C)** The Gubaphage clade was found in 5 continents, with Europe harbouring the highest number of infected samples (38%), as opposed to South America with none detected.

Given the large genetic diversity contained in Gubaphage’s VC we sought to characterize its phylogenetic structure (Figure 5B). Analysis of protein overlap between Gubaphage’s genomes revealed that this clade is composed of 2 clusters that share more than 20% but less than 40% of homologous proteins between them. This structure suggests two genera (G1 and G2) from a single viral subfamily. In addition, within G1 we identified another phylogenetic substructure composed of 3 large clusters (G1.1, G1.2, and G1.3) composed of 313, 514, and 502 phage genomes respectively. Host range prediction revealed that G1.1 infects *Bacteroides caccae* and *Bacteroides xylanisolvens* B, G1.3 *Bacteroides* B *vulgatus*, and G2 *Parabacteroides merdae* and *Parabacteroides distasonis*. In the case of G1.2 we couldn’t confidently predict a putative host. Interestingly, the larger genetic distance between G1 and G2 also resulted in a more extreme host range switch, from Bacteroidaceae (G1) to Porphyromonadaceae (G2). Core genes of the Gubaphage included homing endonucleases, DNA polymerase I, FluMu terminase, DNA primase, DNA helicase, Thymidylate kinase, dUTPase, among others. Annotation of its genome revealed that Gubaphage is organized into three distinct regions. One region encodes structural proteins, the second is composed mainly of genes involved in DNA processing and the third codes for a series of hypothetical proteins. We also found that the phage FAKO05_000032F (Suzuki et al., 2019), had high sequence identity (>90%) with several members of G1.3.

Analysis of the distribution of the Gubaphage clade revealed its presence in all the continents except in South America (Figure 5C). Particularly, it reached a prevalence close to 40% in Europe, while the lowest corresponded to Africa (3%). The discovery of the Gubaphage clade is yet another example of a highly prevalent phage in the human gut and highlights the need to perform further culturing and mechanistic studies to better understand its role in the human gut microbiota.

## DISCUSSION

In this study, we generated and analysed a collection of ∼142,000 high-quality and non-redundant gut phage genomes recovered from 28,060 worldwide distributed human gut metagenomes and 2898 gut isolate genomes. To our knowledge, this set represents the most comprehensive and complete collection of human gut phage genomes to date and is complemented by other published gut phage databases (Gregory et al., 2019; Paez-Espino et al., 2019). Importantly, this work shows that it is possible to recover high-quality phage genomes from shotgun metagenomes without the need to enrich for viral-like particles (VLPs) from stool samples prior to sequencing. With our approach, we not only recovered non-integrative phages like p-crAssphage, but also uncovered prophage sequences which may rarely enter the lytic cycle and form VLPs. As shotgun metagenomes are far more readily available than VLP metagenomes, we had access to an unparalleled number of datasets which enabled us to obtain more complete genomes and viral diversity. Our pipeline highlighted the need for stringent quality control procedures in order to filter out contamination when dealing with predictions of mobile genetic elements such as phages. This is particularly true when mining large-scale datasets due to the impossibility of manually curating every prediction. As the field moves towards the analysis of larger datasets, we believe that machine learning approaches (such as the classifier developed in this work) can be harnessed to help mitigate contamination and significantly boost the quality of the final set of predictions.

Grouping our predictions into VCs was a critical aspect to organize and manage the vast number of predictions in our database. VCs allowed us to discover important phageome patterns such as uncovering highly genetically diverse phage clades (p-crAssphage and Gubaphage), inferring host range, evaluating prevalence around the world, and exposing epidemiology associations by profiling the phageome composition of human samples. Although vContact (Bin Jang et al., 2019) has been extensively used to group phage sequences into clusters that roughly correspond to genus level, it was not computationally feasible to use it with our massive database. We foresee that as genomic and phenotypic features of these VCs are further studied, it will be possible to classify them into at least one of the 15 hierarchical ranks recommended by the ICTV.

Here we also carried out the most comprehensive analysis of the host range of human gut phages. Although the majority of VCs were found to be restricted to a single bacterial species, a significant percentage (∼36%) was predicted to infect multiple species, genera, families, orders, and even classes. A consequence of broad host range phages is an increased connectivity for horizontal gene transfer events due to transduction, which can result in the creation of gene flow networks between phylogenetically distinct bacterial species in the human gut.

The use of GPD also enabled us to gain new insights into the epidemiology of gut phages. Notably, we were able to harness global variation in phage composition to learn that the human gut phageome is associated with the lifestyle of individuals and populations. We showed that phages found in urban samples targeted *Bacteroides* over Prevotellaceae bacteria, whereas rural samples from Peru, Tanzania, Madagascar, and Fiji harboured phages with a host range that targeted Prevotellaceae over *Bacteroides* bacteria. This is yet another result that highlights the importance of the size and diversity of our initial dataset, as we were able to capture the genomes of phages from several understudied regions.

In this work, we also show how our newly generated GPD can be harnessed for characterization of other important viral subfamilies from the gut. In particular, we discovered that the novel Gubaphage clade was actually composed of 2 genera and was able to infect bacteria from the Bacteroidaceae and Porphyromonadaceae families. The combined prevalence of the 2 Gubaphage genera reached a sample proportion between 10-15% in North America, Oceania and Asia, while in Europe it was found to be infecting bacteria in ∼37% of the samples. These results highlight the importance of establishing well-defined viral gut subfamilies, as the combined effect size of highly related phage genomes may help uncover associations of specific clades with their bacterial hosts and human health.

Having a comprehensive database of high-quality phage genomes paves the way for a multitude of analyses of the human gut virome at a greatly improved resolution, enabling the association of specific viral clades with distinct microbiome phenotypes. Importantly, GPD provides a blueprint to guide functional and phenotypic experiments of the human gut phageome, as we linked over 40,000 predictions to 472 cultured gut bacteria species. GPD also harbours 2496 phages that were mined from cultured isolates that are publicly available, and notably 131 members of 57 globally distributed VCs, providing a resource for wet lab experiments to study bacteria-phage co-evolution. In addition, having more complete genomes allows inspection of the most amenable phages for genetic engineering (Chen et al., 2017) or identification of the receptor binding protein genes to expand their host range (Yehl et al., 2019). Given how important the mobilome can be for bacterial ecosystems, we believe that further characterization of other prominent genetic elements such as ICEs, IMEs, genetic islands, and transposons will bring us closer to understanding the association of the gut microbiome with different lifestyles, age and ultimately, health and disease.

## METHODS

### Metagenome assembly

Sequencing reads from 28,060 human gut metagenomes were obtained from the European Nucleotide Archive (Leinonen et al., 2011) Paired-end reads were assembled using SPAdes v3.10.0 (Bankevich et al., 2012) with option ‘--meta’, while single-end reads were assembled with MEGAHIT v1.1.3 (Li et al., 2015) both with default parameters.

### Viral sequence prediction

To identify viral sequences among human gut metagenomes, we used virFinder (Ren et al., 2017) which relies on k-mer signatures to discriminate viral from bacterial contigs, and VirSorter (Roux et al., 2015) which exploits sequence similarity to known phage and other viral-like features such as GC skew. While VirFinder is only able to classify whole contigs, VirSorter can also detect prophages and thus classifies viral sequences as ‘free’ or integrated. Since obtaining high-quality genomes was paramount for our downstream analyses, we used conservative settings for both tools. Metagenome assembled contigs >10 kb in length were analysed with VirSorter v1.0.5 and VirFinder v1.1 to detect putative viral sequences. With VirSorter, only predictions classified as category 1, 2, 4 or 5 were considered. In the case of VirFinder, we selected contigs with a score >0.9 and *P* <0.01.

Contigs were further quality-filtered to remove host sequences using a blast-based approach. Briefly, we first used the ‘blastn’ function of BLAST v2.6.0 (Altschul et al., 1990) to query each contig against the human genome GRCh38 using the following parameters: ‘-word_size 28 -best_hit_overhang 0.1 - best_hit_score_edge 0.1 -dust yes -evalue 0.0001 -min_raw_gapped_score 100 -penalty −5 - perc_identity 90 -soft_masking true’. Contigs with positive hits across >60% total length were excluded.

### Sequence clustering

Dereplication of the filtered contigs was performed with CD-HIT v4.7 (Li and Godzik, 2006) using a global identity threshold of 99% (‘-c 0.99’). This was performed first on contigs obtained within the same ENA study, and afterwards among those obtained across studies. A final set of representative viral sequences was generated by clustering these resulting contigs at a 95% nucleotide identity over a local alignment of 75% of the shortest sequence (options ‘-c 0.95 -G 0 -aS 0.75’).

### Quality control of GPD predictions

In order to ensure a high-quality of GPD predictions we removed integrative and conjugative elements by using a machine learning approach.

Our training set consisted of all experimental ICEs with intact sequence retrieved from ICEberg 2.0 (Bi et al., 2012) and the phage RefSeq genomes from NCBI (Brister et al., 2015). Our test set was downloaded from the Intestinal microbiome mobile elements database (ImmeDB) corresponding to the “ICEs” and “Prophages” datasets. By parsing GFF files with custom Python scripts, for each sequence we calculated 3 high-level features, namely number of genes/kb, number of hypothetical proteins/total genes, and 5-kmer relative frequency (4^5^ = 1024 kmers). We used Keras with the TensorFlow (Abadi et al.) backend to train a feedforward neural network with an initial hidden layer of size 10 (ReLU activation), followed by another hidden layer of size 5 (ReLU activation) and a final neuron with a sigmoid activation function. Model selection was carried out with 5-fold cross-validation. We trained the network using the Adam optimizer and the binary cross entropy as the loss function.

We carried out the classification by allowing a false positive rate of 0.25% with a recall of 91%. Finally, we excluded genomes that were predicted to belong to non-phage taxa (82 predictions)

### Clustering of phages into VCs

We first created a BLAST database (makeblastdb) of all the nucleotide sequences stored in GPD and then carried out all the pairwise comparisons by blasting GPD against itself (we kept hits with E-value ≤0.001). Then, for every pairwise comparison, we calculated the coverage by merging the aligned fraction length of the smaller sequence that shared at least 90% sequence similarity. We kept only the results with a coverage >75%. Finally, we carried out a graph-based clustering by running the Markov Clustering Algorithm (MCL) (Dongen, 2000) with an inflation value of 6.0.

### Viral taxonomic assignment

Viral taxonomic assignment of contigs was performed using a custom database of phylogenetically informative profile HMMs (ViPhOG v1, available here: ftp://ftp.ebi.ac.uk/pub/databases/metagenomics/viral-pipeline/hmmer_databases), where each model is specific to one viral taxon. First, protein-coding sequences of each viral contig were predicted with Prodigal v2.6.3 (Hyatt et al., 2010). Thereafter, we used ‘hmmscan’ from HMMER v3.1b2 (Eddy, 1998) to query each protein sequence against the ViPhOG database, setting a full-sequence E-value reporting threshold of 10^−3^ and a per-domain independent E-value threshold of 0.1. Resulting hits were analysed to predict the most likely and specific taxon for the whole contig based on the following criteria: (i) a minimum of 20% of genes with hits against the ViPhOG database, or at least two genes if the contig had less than 10 total genes; and (ii) among those with hits against the ViPhOG database, a minimum of 60% assigned to the same viral taxon.

### Metagenomic read mapping

To estimate the prevalence of each viral species, we mapped metagenomic reads using BWA-MEM v0.7.16a-r1181 (Li and Durbin, 2009) (‘bwa mem -M’) against the GPD database (clustered at 95% nucleotide identity) here generated. Mapped reads were filtered with samtools v1.5 (Li et al., 2009) to remove secondary alignments (‘samtools view -F 256’) and each viral species was considered present in a sample if the mapped reads covered >75% of the genome length.

### Taxonomic assignment of bacterial genomes

Bacterial isolate genomes were taxonomically classified with the Genome Taxonomy Database Toolkit (GTDB-Tk) v0.3.1 (Chaumeil et al., 2019) (https://github.com/Ecogenomics/GTDBTk) (database release 04-RS89) using the ‘classify_wf’ function and default parameters. Taxa with an alphabetic suffix represent lineages that are polyphyletic or were subdivided due to taxonomic rank normalization according to the GTDB reference tree. The unsuffixed lineage contains the type strain whereas all other lineages are given alphabetic suffixes, suggesting that their labelling should be revised in due course.

### Clustering of proteins

We predicted the whole proteome of GPD with Prodigal v2.6.3 (metagenomic mode) and masked the low-complexity regions with DustMasker. We then created a BLAST database of all the protein sequences and carried out all the pairwise comparisons by blasting the GPD proteome against itself (we kept hits with E-value ≤0.001). Then, for every pairwise comparison, we calculated a similarity metric as defined by Chan et al (Chan et al., 2013). Finally, we ran the Markov Clustering Algorithm (MCL) with an inflation value of 6.0 and removed clusters with only 1 member.

### Geographical distribution of metagenomic samples

We removed samples with a sequencing depth below 50 million reads/sample, as below this threshold we observed a positive correlation between sample depth and number of viral genomes detected (Supplemental figure 3B). This new subset consisted of 3011 samples and spanned all the continents and 23 countries. Similarity between 2 samples was calculated by computing the number of shared VCs divided by the total number of VCs in both samples (Jaccard index). Distribution of samples was visualized with PCA.

### Host assignment

We predicted CRISPR spacer sequences from the 2898 gut bacteria using CrisprCasFinder-2.0.2 (Couvin et al., 2018). We only used spacers found in CRISPR arrays having evidence levels 3 and 4. We assigned a host to a prediction only if the putative host CRISPR spacer matched perfectly to the phage prediction (100% sequence identity across whole length of CRISPR spacer). We carried out the screen by blasting all the predicted CRISPR spacers against the nucleotide GPD BLAST database using the following custom settings (task: blastn-short, - gapopen 10, -gapextend 2, penalty −1, - word_size 7m -perc_identity 100). We kept only hits that matched across the whole length of the spacer with a custom script. In addition, prophages were assigned to the bacterial assembly from which they were predicted.

### Phylogenetic analysis of Gubaphage

The phylogenetic tree comparing Gubaphage against crAss-like phages was constructed by aligning the corresponding large terminase genes with MAFFT v7.453 (Katoh et al., 2002) –auto mode, followed by FastTree v2.1.10 (Price et al., 2010). The resultant tree was visualized on iTOL (Letunic and Bork, 2007). We calculated the fraction of shared protein clusters among all the Gubaphage genomes and then carried out hierarchical clustering with average linkage and Euclidean metric.

### Annotation of viral genomes

Protein annotation was carried out using Prokka v1.5-135 (Seemann, 2014).

## Data and code availability

GPD sequences and associated metadata can be found in the following FTP link: http://ftp.ebi.ac.uk/pub/databases/metagenomics/genome_sets/gut_phage_database/Classifier and scripts used to generate figures can be found here: https://github.com/cai91/GPD

## Acknowledgements

This work was supported by the Wellcome Trust (098051). L.F.C. is supported by a Wellcome Sanger Institute PhD Studentship. A.A. and R.D.F are funded by EMBL core funds. We would like to thank Y. Shao and H. Browne for fruitful discussions and feedback about the manuscript.

## Author contributions

L.F.C., A.A., R.D.F., and T.D.L. conceived the study. L.F.C. wrote the manuscript and made the figures, assessed quality of GPD predictions, developed the classifier to distinguish phages from ICEs, analysed viral diversity patterns across gut isolates, analysed global epidemiology trends, and defined the Gubaphage clade. A.A. assembled human gut metagenomes, carried out viral prediction and mapped predictions to metagenomes. G.R.P. wrote the phage taxonomic classification pipeline. All authors read, edited and approved the final manuscript.

## Competing interests

T.D.L. and R.D.F. are either employees of, or consultants to, Microbiotica Pty Ltd.

## Supplementary figures

**Figure S1.**
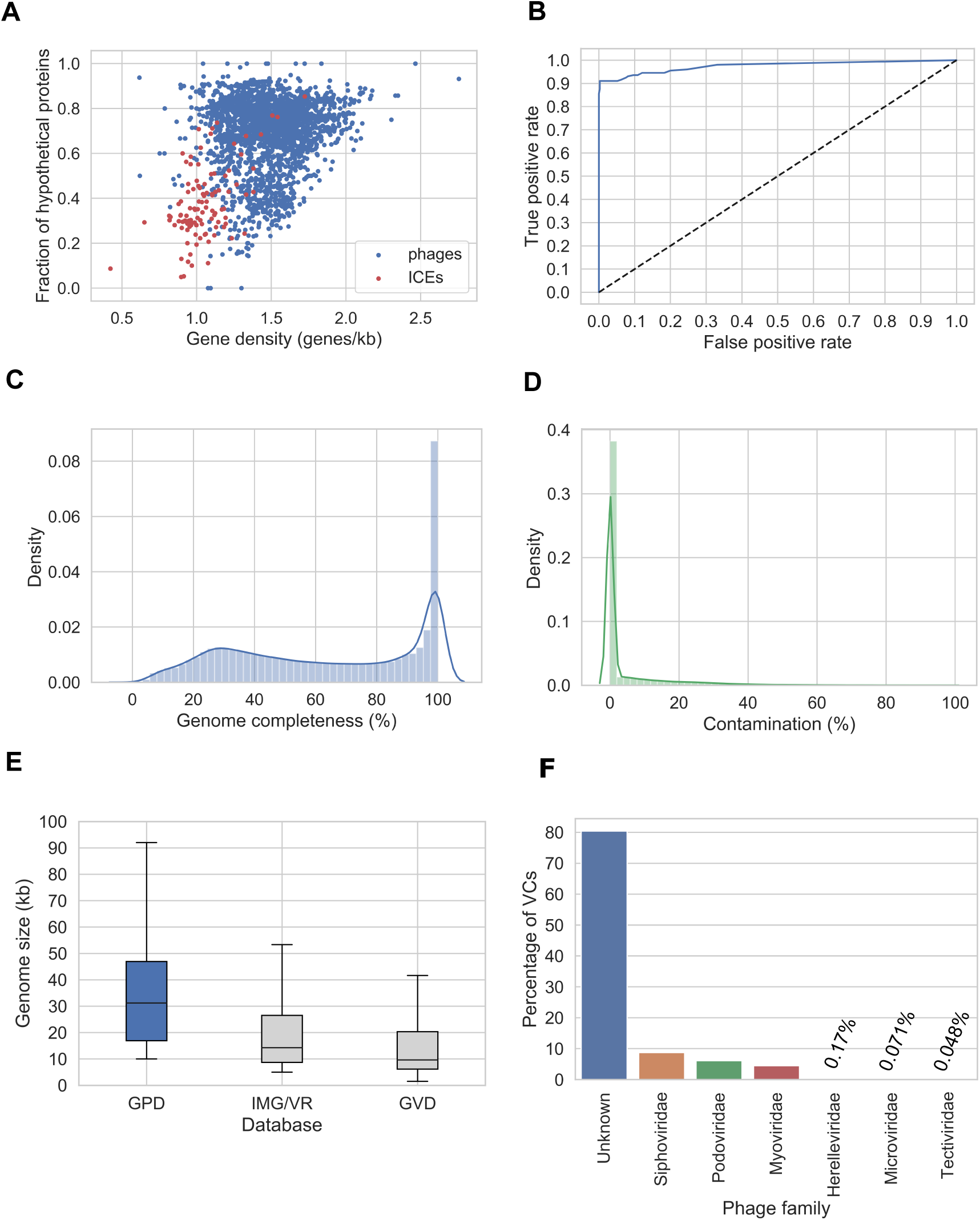
Generating the most complete sequence database of human gut bacteriophages. **A)** Gene density and fraction of hypothetical proteins are features that can be harnessed discriminate phages from ICEs. **B)** ROC curve showing the high performance (AUC>0.97) of the neural network developed to decontaminate ICEs from phages. **C**) Genome completeness distribution as estimated by CheckV on GPD. D) GPD contamination distribution according to CheckV. **D)** Size distribution of GPD against other public databases. **E)** Assignment of viral taxonomy to GPD predictions.

**Figure S2.**
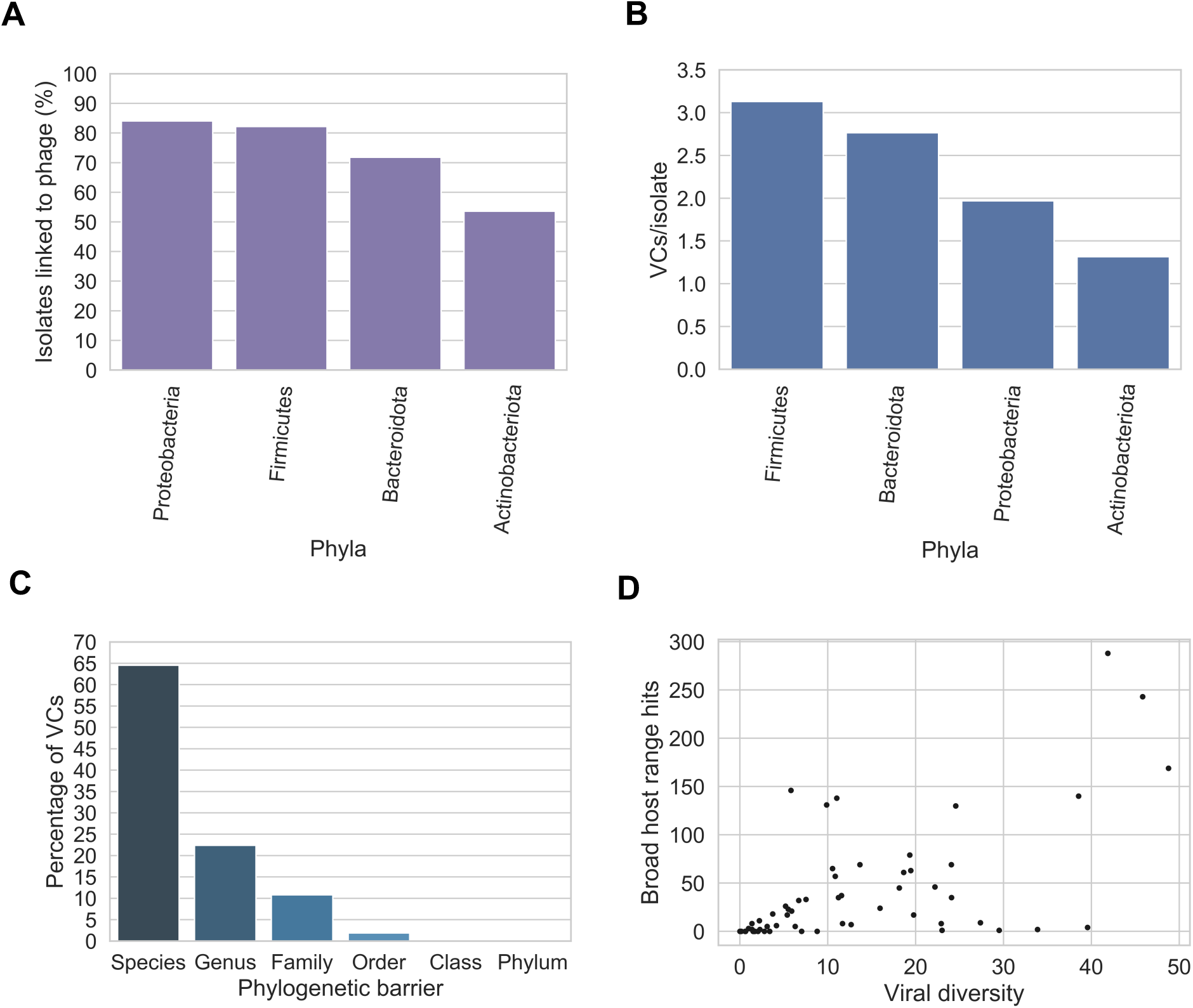
Bacterial host assignment and host range for gut phage. **A)** Percentage of isolates of each phylum linked to phage by CRISPR spacers and prophage assignment. Actinobacteria had the lowest percentage of isolates predicted to be a phage host. Actinobacteria vs Bacteroidota (*P =* 0.007, *χ*^2^ test), Actinobacteria vs Proteobacteria (*P =* 0.0025, *χ*^2^ test), Actinobacteria vs Firmicutes (*P =* 1.01⨯ 10^−5^, *χ*^2^ test). **B)** The Firmicutes hosted the highest viral diversity (highest number of VCs/isolate). Firmicutes vs Bacteroidota (*P =* 0.021, *χ*^2^ test), Firmicutes vs Proteobacteria (*P =* 4.41 ⨯ 10^−6,^ *χ*^2^ test), Firmicutes vs Actinobacteriota (*P =* 1.1 ⨯ 10^−31^, *χ*^2^ test) **C)** The majority of VCs were found to be restricted to infect a single species. However, a considerable number of VCs (∼36%) had a broader host range (*P =* 0.0, binomial test). **D)** In general, the higher the viral diversity per bacterial genus, the higher the number of phages with broad host range (Spearman’s Rho = 0.6685, *P =* 3.91⨯ 10-9).

**Figure S3.**
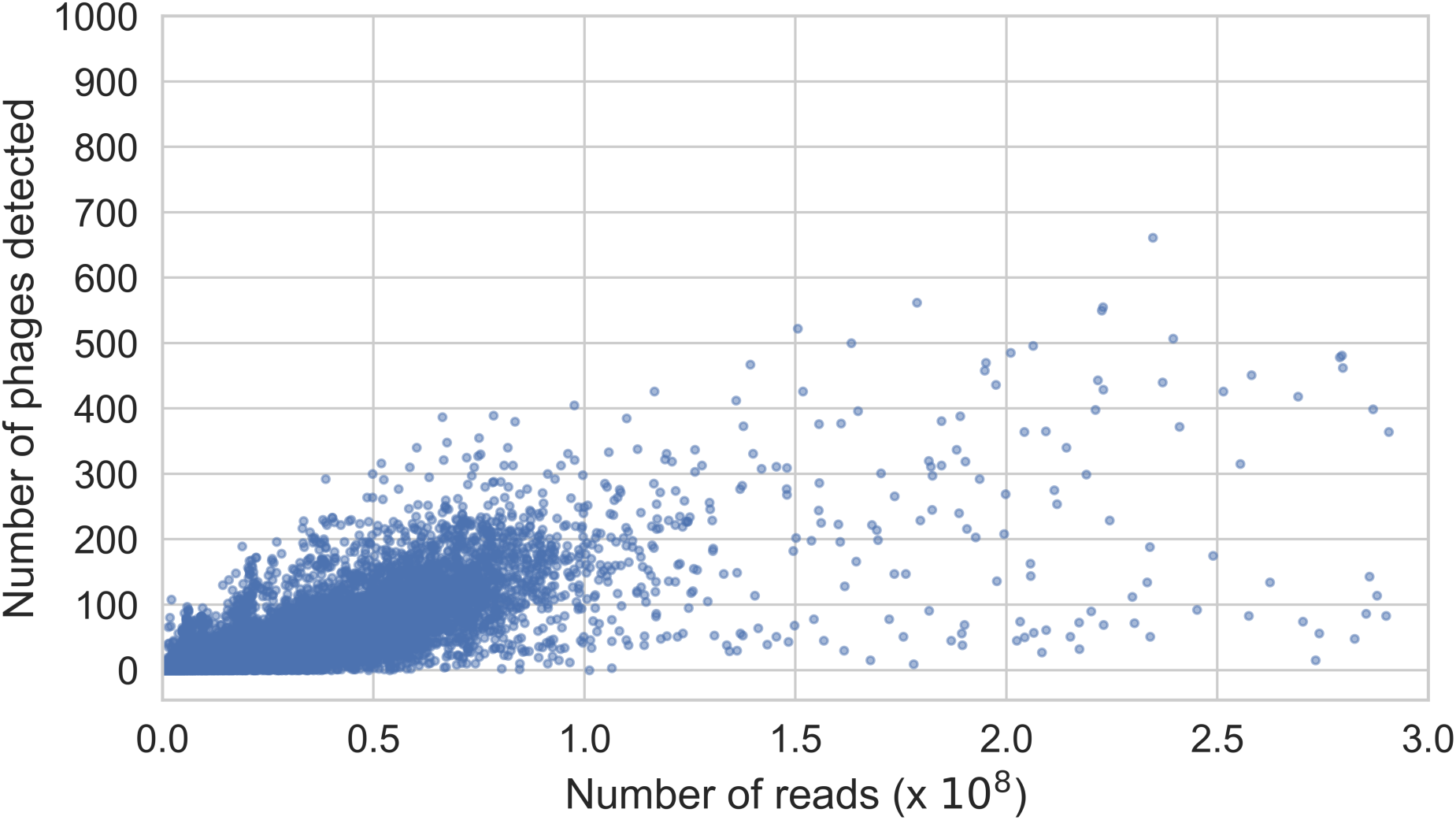
Relationship between sample sequencing depth and phage richness. Samples exhibit a positive correlation between sequencing depth and number of phage genomes detected. In order to reduce this bias, we analysed only samples with a sequencing depth >50 million reads/sample. Correlation of samples with sequencing depth <50 million (Pearson’s r: 0.6825, *P =* 0.0). Correlation of samples with sequencing depth >50 million (Pearson’s r: 0.3681, *P =* 2.79e-97).

**Figure S4.**
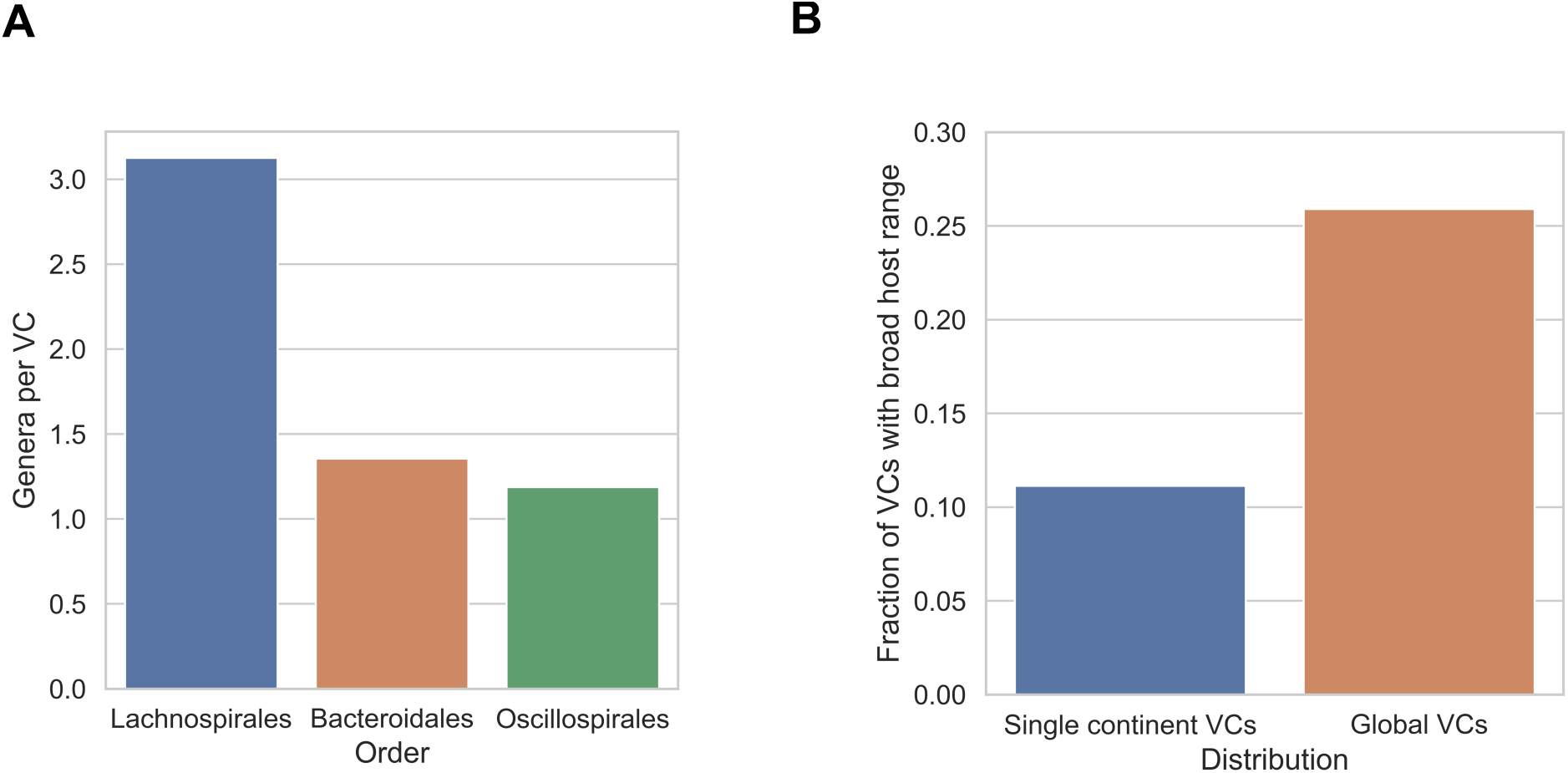
Global gut phage clades and their bacterial hosts. **A)** When analysing globally distributed VCs, the VCs from the order of Lachnospirales were shared across a wider range of genera than those within Oscillospirales and Bacteroidales. Lachnospirales vs Bacteroidales (*P =* 9.99 ⨯ 10^−6^, *χ*^2^ test). Lachnospirales vs Oscillospirales (*P =* 6.55 ⨯ 10^−6^, *χ*^2^ test). **B)** We observed that globally distributed phages had a significantly broader range (above genus) than phages found in single continents (*P =* 1.63 ⨯ 10^−^ 5, *χ*^2^ test).

**Figure S5.**
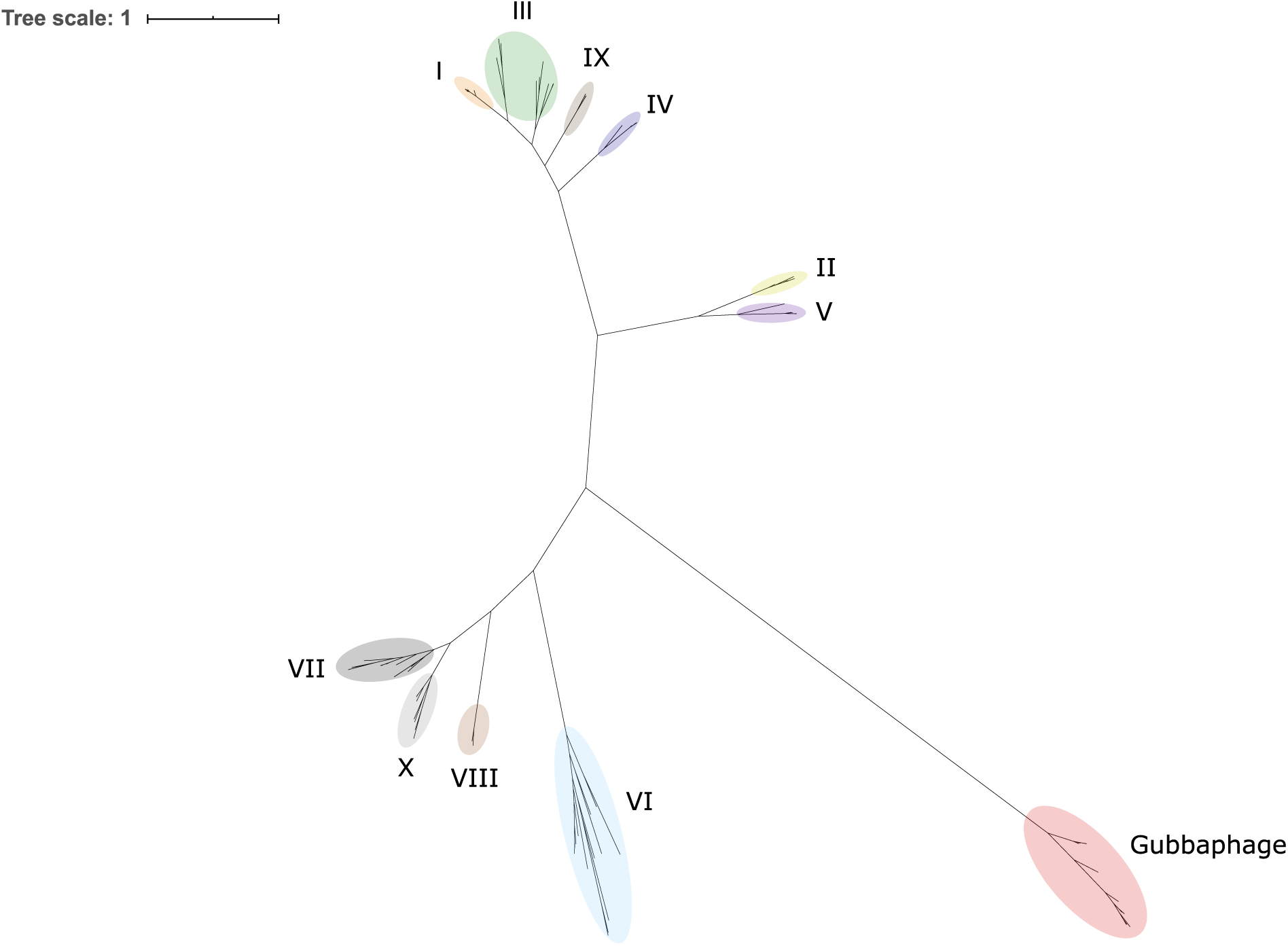
The Gubaphage is a novel and highly prevalent clade in the human gut. Unrooted phylogenetic tree of the large terminase gene from 226 crAss-like genomes and 44 Gubaphage sequences. Roman numerals correspond to the 10 crass-like genera. The Gubaphage significantly diverged from other crAss-like phages forming a distant clade of its own (red).

## Notes

http://ftp.ebi.ac.uk/pub/databases/metagenomics/genome_sets/gut_phage_database/

https://github.com/cai91/GPD

## REFERENCES

Abadi, M., Barham, P., Chen, J., Chen, Z., Davis, A., Dean, J., Devin, M., Ghemawat, S., Irving, G., Isard, M., et al. TensorFlow: A system for large-scale machine learning. 21.

Ackermann, H.W. (1998). Tailed bacteriophages: the order caudovirales. Adv. Virus Res. 51, 135–201.

Al-Shayeb, B., Sachdeva, R., Chen, L.-X., Ward, F., Munk, P., Devoto, A., Castelle, C.J., Olm, M.R., Bouma-Gregson, K., Amano, Y., et al. (2020). Clades of huge phages from across Earth’s ecosystems. Nature 578, 425–431.

Altschul, S.F., Gish, W., Miller, W., Myers, E.W., and Lipman, D.J. (1990). Basic local alignment search tool. J. Mol. Biol. 215, 403–410.

Bankevich, A., Nurk, S., Antipov, D., Gurevich, A.A., Dvorkin, M., Kulikov, A.S., Lesin, V.M., Nikolenko, S.I., Pham, S., Prjibelski, A.D., et al. (2012). SPAdes: A New Genome Assembly Algorithm and Its Applications to Single-Cell Sequencing. J. Comput. Biol. 19, 455–477.

Barr, J.J., Auro, R., Furlan, M., Whiteson, K.L., Erb, M.L., Pogliano, J., Stotland, A., Wolkowicz, R., Cutting, A.S., Doran, K.S., et al. (2013). Bacteriophage adhering to mucus provide a non-host-derived immunity. Proc. Natl. Acad. Sci. U. S. A. 110, 10771–10776.

Betts, A., Kaltz, O., and Hochberg, M.E. (2014). Contrasted coevolutionary dynamics between a bacterial pathogen and its bacteriophages. Proc. Natl. Acad. Sci. U. S. A. 111, 11109–11114.

Bi, D., Xu, Z., Harrison, E.M., Tai, C., Wei, Y., He, X., Jia, S., Deng, Z., Rajakumar, K., and Ou, H.-Y. (2012). ICEberg: a web-based resource for integrative and conjugative elements found in Bacteria. Nucleic Acids Res. 40, D621–D626.

Bin Jang, H., Bolduc, B., Zablocki, O., Kuhn, J.H., Roux, S., Adriaenssens, E.M., Brister, J.R., Kropinski, A.M., Krupovic, M., Lavigne, R., et al. (2019). Taxonomic assignment of uncultivated prokaryotic virus genomes is enabled by gene-sharing networks. Nat. Biotechnol. 37, 632–639.

Breitbart, M., Bonnain, C., Malki, K., and Sawaya, N.A. (2018). Phage puppet masters of the marine microbial realm. Nat. Microbiol. 3, 754–766.

Brister, J.R., Ako-adjei, D., Bao, Y., and Blinkova, O. (2015). NCBI Viral Genomes Resource. Nucleic Acids Res. 43, D571–D577.

Brüssow, H., and Hendrix, R.W. (2002). Phage genomics: small is beautiful. Cell 108, 13–16.

Chan, C.X., Mahbob, M., and Ragan, M.A. (2013). Clustering evolving proteins into homologous families. BMC Bioinformatics 14, 120.

Chaumeil, P.-A., Mussig, A.J., Hugenholtz, P., and Parks, D.H. (2019). GTDB-Tk: a toolkit to classify genomes with the Genome Taxonomy Database. Bioinforma. Oxf. Engl.

Chen, J., Quiles-Puchalt, N., Chiang, Y.N., Bacigalupe, R., Fillol-Salom, A., Chee, M.S.J., Fitzgerald, J.R., and Penadés, J.R. (2018). Genome hypermobility by lateral transduction. Science 362, 207–212.

Chen, M., Zhang, L., Abdelgader, S.A., Yu, L., Xu, J., Yao, H., Lu, C., and Zhang, W. (2017). Alterations in gp37 Expand the Host Range of a T4-Like Phage. Appl. Environ. Microbiol. 83.

Clooney, A.G., Sutton, T.D.S., Shkoporov, A.N., Holohan, R.K., Daly, K.M., O’Regan, O., Ryan, F.J., Draper, L.A., Plevy, S.E., Ross, R.P., et al. (2019). Whole-Virome Analysis Sheds Light on Viral Dark Matter in Inflammatory Bowel Disease. Cell Host Microbe 26, 764–778.e5.

Couvin, D., Bernheim, A., Toffano-Nioche, C., Touchon, M., Michalik, J., Néron, B., Rocha, E.P.C., Vergnaud, G., Gautheret, D., and Pourcel, C. (2018). CRISPRCasFinder, an update of CRISRFinder, includes a portable version, enhanced performance and integrates search for Cas proteins. Nucleic Acids Res. 46, W246–W251.

Dongen, S.M. van (2000). Graph clustering by flow simulation.

Dutilh, B.E., Cassman, N., McNair, K., Sanchez, S.E., Silva, G.G.Z., Boling, L., Barr, J.J., Speth, D.R., Seguritan, V., Aziz, R.K., et al. (2014). A highly abundant bacteriophage discovered in the unknown sequences of human faecal metagenomes. Nat. Commun. 5, 4498.

Eddy, S.R. (1998). Profile hidden Markov models. Bioinforma. Oxf. Engl. 14, 755–763.

Edwards, R.A., McNair, K., Faust, K., Raes, J., and Dutilh, B.E. (2016). Computational approaches to predict bacteriophage-host relationships. FEMS Microbiol. Rev. 40, 258–272.

Forster, S.C., Kumar, N., Anonye, B.O., Almeida, A., Viciani, E., Stares, M.D., Dunn, M., Mkandawire, T.T., Zhu, A., Shao, Y., et al. (2019). A human gut bacterial genome and culture collection for improved metagenomic analyses. Nat. Biotechnol. 37, 186–192.

Gregory, A.C., Zablocki, O., Howell, A., Bolduc, B., and Sullivan, M.B. (2019). The human gut virome database. BioRxiv 655910.

Guerin, E., Shkoporov, A., Stockdale, S.R., Clooney, A.G., Ryan, F.J., Sutton, T.D.S., Draper, L.A., Gonzalez-Tortuero, E., Ross, R.P., and Hill, C. (2018). Biology and Taxonomy of crAss-like Bacteriophages, the Most Abundant Virus in the Human Gut. Cell Host Microbe 24, 653–664.e6.

Harrison, E., and Brockhurst, M.A. (2017). Ecological and Evolutionary Benefits of Temperate Phage: What Does or Doesn’t Kill You Makes You Stronger. BioEssays 39, 1700112.

Hoyles, L., McCartney, A.L., Neve, H., Gibson, G.R., Sanderson, J.D., Heller, K.J., and van Sinderen, D. (2014). Characterization of virus-like particles associated with the human faecal and caecal microbiota. Res. Microbiol. 165, 803–812.

Hyatt, D., Chen, G.-L., Locascio, P.F., Land, M.L., Larimer, F.W., and Hauser, L.J. (2010). Prodigal: prokaryotic gene recognition and translation initiation site identification. BMC Bioinformatics 11, 119.

Jahn, M.T., Arkhipova, K., Markert, S.M., Stigloher, C., Lachnit, T., Pita, L., Kupczok, A., Ribes, M., Stengel, S.T., Rosenstiel, P., et al. (2019). A Phage Protein Aids Bacterial Symbionts in Eukaryote Immune Evasion. Cell Host Microbe 26, 542–550.e5.

Jain, R., Rivera, M.C., and Lake, J.A. (1999). Horizontal gene transfer among genomes: the complexity hypothesis. Proc. Natl. Acad. Sci. U. S. A. 96, 3801–3806.

Katoh, K., Misawa, K., Kuma, K., and Miyata, T. (2002). MAFFT: a novel method for rapid multiple sequence alignment based on fast Fourier transform. Nucleic Acids Res. 30, 3059–3066.

Kho, Z.Y., and Lal, S.K. (2018). The Human Gut Microbiome - A Potential Controller of Wellness and Disease. Front. Microbiol. 9, 1835.

Koert, M., Mattson, C., Caruso, S., and Erill, I. (2019). Evidence for shared ancestry between Actinobacteria and Firmicutes bacteriophages. BioRxiv 842583.

Leinonen, R., Akhtar, R., Birney, E., Bower, L., Cerdeno-Tárraga, A., Cheng, Y., Cleland, I., Faruque, N., Goodgame, N., Gibson, R., et al. (2011). The European Nucleotide Archive. Nucleic Acids Res. 39, D28–D31.

Letunic, I., and Bork, P. (2007). Interactive Tree Of Life (iTOL): an online tool for phylogenetic tree display and annotation. Bioinforma. Oxf. Engl. 23, 127–128.

Li, H., and Durbin, R. (2009). Fast and accurate short read alignment with Burrows–Wheeler transform. Bioinformatics 25, 1754–1760.

Li, W., and Godzik, A. (2006). Cd-hit: a fast program for clustering and comparing large sets of protein or nucleotide sequences. Bioinforma. Oxf. Engl. 22, 1658–1659.

Li, D., Liu, C.-M., Luo, R., Sadakane, K., and Lam, T.-W. (2015). MEGAHIT: an ultra-fast single-node solution for large and complex metagenomics assembly via succinct de Bruijn graph. Bioinformatics 31, 1674–1676.

Li, H., Handsaker, B., Wysoker, A., Fennell, T., Ruan, J., Homer, N., Marth, G., Abecasis, G., Durbin, R., and 1000 Genome Project Data Processing Subgroup (2009). The Sequence Alignment/Map format and SAMtools. Bioinforma. Oxf. Engl. 25, 2078–2079.

Manrique, P., Bolduc, B., Walk, S.T., Oost, J. van der, Vos, W.M. de, and Young, M.J. (2016). Healthy human gut phageome. Proc. Natl. Acad. Sci. 113, 10400–10405.

Marbouty, M., Thierry, A., and Koszul, R. (2020). Phages - bacteria interactions network of the healthy human gut (Microbiology).

Minot, S., Grunberg, S., Wu, G.D., Lewis, J.D., and Bushman, F.D. (2012). Hypervariable loci in the human gut virome. Proc. Natl. Acad. Sci. U. S. A. 109, 3962–3966.

Nayfach, S., Camargo, A.P., Eloe-Fadrosh, E., Roux, S., and Kyrpides, N. (2020). CheckV: assessing the quality of metagenome-assembled viral genomes. BioRxiv 2020.05.06.081778.

Paez-Espino, D., Eloe-Fadrosh, E.A., Pavlopoulos, G.A., Thomas, A.D., Huntemann, M., Mikhailova, N., Rubin, E., Ivanova, N.N., and Kyrpides, N.C. (2016). Uncovering Earth’s virome. Nature 536, 425–430.

Paez-Espino, D., Roux, S., Chen, I.-M.A., Palaniappan, K., Ratner, A., Chu, K., Huntemann, M., Reddy, T.B.K., Pons, J.C., Llabrés, M., et al. (2019). IMG/VR v.2.0: an integrated data management and analysis system for cultivated and environmental viral genomes. Nucleic Acids Res. 47, D678–D686.

Price, M.N., Dehal, P.S., and Arkin, A.P. (2010). FastTree 2 – Approximately Maximum-Likelihood Trees for Large Alignments. PLoS ONE 5.

Ren, J., Ahlgren, N.A., Lu, Y.Y., Fuhrman, J.A., and Sun, F. (2017). VirFinder: a novel k-mer based tool for identifying viral sequences from assembled metagenomic data. Microbiome 5, 69.

Reyes, A., Haynes, M., Hanson, N., Angly, F.E., Heath, A.C., Rohwer, F., and Gordon, J.I. (2010). Viruses in the fecal microbiota of monozygotic twins and their mothers. Nature 466, 334–338.

Roux, S., Krupovic, M., Poulet, A., Debroas, D., and Enault, F. (2012). Evolution and Diversity of the Microviridae Viral Family through a Collection of 81 New Complete Genomes Assembled from Virome Reads. PLoS ONE 7.

Roux, S., Enault, F., Hurwitz, B.L., and Sullivan, M.B. (2015). VirSorter: mining viral signal from microbial genomic data. PeerJ 3.

Roux, S., Adriaenssens, E.M., Dutilh, B.E., Koonin, E.V., Kropinski, A.M., Krupovic, M., Kuhn, J.H., Lavigne, R., Brister, J.R., Varsani, A., et al. (2019). Minimum Information about an Uncultivated Virus Genome (MIUViG).Nat. Biotechnol. 37, 29–37.

Seemann, T. (2014). Prokka: rapid prokaryotic genome annotation. Bioinforma. Oxf. Engl. 30, 2068–2069.

Simmonds, P., Adams, M.J., Benkő, M., Breitbart, M., Brister, J.R., Carstens, E.B., Davison, A.J., Delwart, E., Gorbalenya, A.E., Harrach, B., et al. (2017). Consensus statement: Virus taxonomy in the age of metagenomics. Nat. Rev. Microbiol. 15, 161–168.

Suzuki, Y., Nishijima, S., Furuta, Y., Yoshimura, J., Suda, W., Oshima, K., Hattori, M., and Morishita, S. (2019). Long-read metagenomic exploration of extrachromosomal mobile genetic elements in the human gut. Microbiome 7.

Wu, G.D., Chen, J., Hoffmann, C., Bittinger, K., Chen, Y.-Y., Keilbaugh, S.A., Bewtra, M., Knights, D., Walters, W.A., Knight, R., et al. (2011). Linking Long-Term Dietary Patterns with Gut Microbial Enterotypes. Science 334, 105–108.

Yehl, K., Lemire, S., Yang, A.C., Ando, H., Mimee, M., Torres, M.D.T., de la Fuente-Nunez, C., and Lu, T.K. (2019). Engineering Phage Host-Range and Suppressing Bacterial Resistance through Phage Tail Fiber Mutagenesis. Cell 179, 459–469.e9.

Zou, Y., Xue, W., Luo, G., Deng, Z., Qin, P., Guo, R., Sun, H., Xia, Y., Liang, S., Dai, Y., et al. (2019). 1,520 reference genomes from cultivated human gut bacteria enable functional microbiome analyses. Nat. Biotechnol. 37, 179–185.

