## Supplementary material for "Massive expansion of human gut bacteriophage diversity": Table S1

| **crAss-like genus** | **Predicted hosts** |
| --- | --- |
| I | NA |
| II | NA |
| III | *Bacteroides*_B *vulgatus* |
| IV | *Bacteroides xylanisolvens*_B |
|  | *Bacteroides caccae* |
| V | *Fusicatenibacter saccharivorans* |
|  | *Lachnospira eligens* |
| VI | *Bacteroides xylanisolvens*_B |
|  | *Prevotella copri* |
| VII | *Bacteroides thetaiotaomicron* |
|  | *Bacteroides*_B *massiliensis* |
|  | *Bacteroides*_B *dorei* |
|  | *Bacteroides caccae* |
|  | *Bacteroides faecis* |
|  | *Bacteroides eggerthii* |
|  | *Bacteroides xylanisolvens*_B |
|  | *Bacteroides*_B *vulgatus* |
|  | *Bacteroides uniformis* |
| VIII | *Prevotella copri* |
| IX | *Prevotella copri* |
| X | *Parabacteroides merdae* |
|  | *Parabacteroides distasonis* |

**Table S1.** Predicted hosts of the crAss-like family
