## Supplementary material for "Massive expansion of human gut bacteriophage diversity": Table S2

| **Global_VC** | **GPD_phage_ID** | **Identifier** | **Source** | **Sample** | **Culture Collection** | **Culture Collection Number** |
| --- | --- | --- | --- | --- | --- | --- |
| VC_2 | ivig_4281 | ERR2230082 | HBC | Bacteroides dorei ERR2230082 | WSI | ERR2230082 |
| VC_2 | ivig_3443 | ERR1022271 | HBC | Bacteroides uniformis ERR1022271 | WSI | ERR1022271 |
| VC_2 | ivig_4319 | ERR2230141 | HBC | Bacteroides uniformis ERR2230141 | WSI | ERR2230141 |
| VC_2 | ivig_4298 | ERR2230119 | HBC | Bacteroides uniformis ERR2230119 | WSI | ERR2230119 |
| VC_2 | ivig_3716 | ERR1022414 | HBC | Bacteroides cellulosilyticus ERR1022414 | DSMZ | DSM 108229 |
| VC_2 | ivig_3679 | ERR2221211 | HBC | Bacteroides acidifaciens ERR2221211 | WSI | ERR2221211 |
| VC_2 | ivig_3715 | ERR1022413 | HBC | Bacteroides thetaiotaomicron ERR1022413 | WSI | ERR1022413 |
| VC_2 | ivig_4318 | ERR2230138 | HBC | Bacteroides uniformis ERR2230138 | WSI | ERR2230138 |
| VC_71 | ivig_3350 | ERR2221142 | HBC | Bifidobacterium pseudocatenulatum ERR2221142 | WSI | ERR2221142 |
| VC_65 | ivig_3375 | ERR2221151 | HBC | Lachnospiraceae nov. ERR2221151 | WSI | ERR2221151 |
| VC_94 | ivig_3609 | ERR1022353 | HBC | Lachnospiraceae nov. ERR1022353 | WSI | ERR1022353 |
| VC_65 | ivig_3761 | ERR1022439 | HBC | Eisenbergiella tayi ERR1022439 | WSI | ERR1022439 |
| VC_16 | ivig_3950 | ERR1203972 | HBC | Lachnospiraceae nov. ERR1203972 | WSI | ERR1203972 |
| VC_50 | ivig_4334 | ERR2230158 | HBC | Bifidobacterium longum ERR2230158 | WSI | ERR2230158 |
| VC_50 | ivig_4156 | ERR2221351 | HBC | Bifidobacterium longum ERR2221351 | NCIMB | NCIMB 15144 |
| VC_55 | ivig_3549 | ERR1022318 | HBC | Eubacterium rectale ERR1022318 | WSI | ERR1022318 |
| VC_75 | ivig_3398 | ERR2221161 | HBC | Lachnospiraceae nov. ERR2221161 | WSI | ERR2221161 |
| VC_78 | ivig_4245 | ERR2221398 | HBC | Escherichia coli ERR2221398 | WSI | ERR2221398 |
| VC_138 | ivig_4044 | ERR2221275 | HBC | Dorea formicigenerans ERR2221275 | WSI | ERR2221275 |
| VC_156 | ivig_3731 | ERR1022423 | HBC | Anaerostipes hadrus ERR1022423 | WSI | ERR1022423 |
| VC_156 | ivig_3760 | ERR1022437 | HBC | Fusicatenibacter saccharivorans ERR1022437 | CCUG & JCM | CCUG 68552 & JCM 31268 |
| VC_168 | ivig_3524 | ERR1022310 | HBC | Ruminococcus gnavus ERR1022310 | DSMZ | DSM 108212 |
| VC_168 | ivig_4348 | ERR171257 | HBC | Ruminococcus gnavus ERR171257 | WSI | ERR171257 |
| VC_175 | ivig_3612 | ERR1022356 | HBC | Bifidobacterium pseudocatenulatum ERR1022356 | WSI | ERR1022356 |
| VC_183 | ivig_3767 | ERR1022442 | HBC | Ruminococcaceae nov. ERR1022442 | WSI | ERR1022442 |
| VC_204 | ivig_3670 | ERR1022391 | HBC | Lachnospira nov. ERR1022391 | BCCM | LMG 29490 |
| VC_288 | ivig_3565 | ERR1022327 | HBC | Faecalibacterium prausnitzii ERR1022327 | WSI | ERR1022327 |
| VC_752 | ivig_3547 | ERR1022317 | HBC | Ruminococcaceae nov. ERR1022317 | WSI | ERR1022317 |
| VC_757 | ivig_4035 | ERR2230078 | HBC | Collinsella aerofaciens ERR2230078 | CCUG | CCUG 73007 |
| VC_1461 | ivig_4025 | ERR2230076 | HBC | Lachnoclostridium nov. ERR2230076 | WSI | ERR2230076 |

**Table S2.** Global VCs found in publicly available gut isolates.
